# Placental Insulin-like Growth Factor 1 Insufficiency Drives Neurodevelopmental Disorder‑Relevant Behavioral Changes with Sex‑Specific Vulnerabilities

**DOI:** 10.64898/2026.01.06.697024

**Authors:** Annemarie J. Carver, Faith M. Fairbairn, Robert J. Taylor, Benjamin W.Q. Hing, Amrita Gajmer, Regan T. Fair, Hanna E. Stevens

## Abstract

Preterm birth, placental insufficiency, and other perinatal adversities lead to the loss of placental support including critical hormones, such as Insulin-like growth factor 1 (IGF1), required for neurodevelopment. Decreased IGF1 and preterm birth are associated with neurodevelopmental disorder risk, including autism spectrum disorder. Whether placental *Igf1* insufficiency drives neurodevelopmental risks is not understood. To understand these mechanisms, placental-targeted CRISPR manipulation in mice was employed to induce placental *Igf1* insufficiency. Subsequently, embryonic forebrain development was assessed sex-specifically to identify structural, cellular, and transcriptomic changes. Postnatal offspring were used to determine neurobehavioral trajectories relevant to neurodevelopmental disorders as assessed through learning, motor, and affective behavioral tasks and neurostereology. Placental *Igf1* insufficiency reduced embryonic forebrain growth, including decreased cell population across males and females. Embryonic forebrain transcriptomics revealed sex-specific alterations. Developmental pathways including insulin-like growth factor receptor signaling, laminin processes, and hormone synthesis were downregulated in male forebrain, driven by autism risk genes, *Reln* and *Lama1*. Altered genes in female forebrain were enriched for autism-risk genes including *Grin2b* and *Dync1h1*. Following these transcriptomic differences, postnatal neurobehavioral trajectories were sex-specific. Male offspring uniquely showed reduced motor learning, increased stereotyped behaviors, altered reversal learning, and reduced forebrain neuronal number. Female offspring displayed opposite behavioral changes as males and few changes in forebrain structure. Assessment of both adult male and female offspring forebrain white matter revealed an increased astrocyte population, a phenotype that appears similar to reactive astrogliosis seen in other models of preterm birth and placental insufficiency. The provision of *Igf1* specifically from placenta is critical for offspring forebrain development. This temporary early deficit has persistent sex-specific neurobehavioral effects. These outcomes have relevance for neurodevelopmental disorder risk and highlight mechanisms that could facilitate intervention development for adverse outcomes after early loss of placental hormone support in perinatal adversity.

## Introduction

Preterm birth increases risk for neurodevelopmental disorders (NDDs), including autism spectrum disorder (ASD) [1-3]. Improved survival rates among preterm infants have coincided with an increased prevalence of these conditions [1,4,5], but mechanisms are not understood. Both preterm birth and NDDs are associated with perinatal adversities involving placental dysfunction and insufficiency [6-8]. Preterm birth causes early loss of placental support, including hormones critical for neurodevelopment [9,10]. This includes Insulin-like growth factor 1 (IGF1), an essential hormone that influences both placental function and neurodevelopment [11-14]. Levels of IGF1 and associated proteins at birth predict growth and neurodevelopment in placental insufficiency and preterm infants as well as birth weight in term babies [15-18].

How IGF1 from placenta specifically influences the brain has been minimally studied. IGF1 is primarily produced by endothelial cells in the placental fetal region [19,20]. Within the placenta, IGF1 promotes hormone and nutrient transport necessary for fetal development via IGF1 receptor (IGF1R) signaling [11]. Within the brain, IGF1 regulates and promotes growth, with high forebrain expression of IGF1R [13,21,22]. IGF1 promotes forebrain cell cycle progression and differentiation across cell types [13,22]. Total *Igf1* knockout in mice reduces adult brain weight and striatal parvalbumin-containing neurons [22,23]. In contrast, total *Igf1* overexpression in mice increases embryonic and juvenile brain weight and cell number in forebrain subregions including the cerebral cortex and caudate-putamen [24]. Placental *Igf1* overexpression increases embryonic striatal growth and alters striatal neurons and striatal-dependent, NDD-relevant behaviors postnatally [13].

Morphological changes to the forebrain and its subregions are associated with NDDs, particularly the striatum which is sensitive to IGF1 changes [13,22,23]. Changes in striatal morphology and function are linked to ASD and ADHD [25-27], including restricted, repetitive behaviors, motor regulation, and procedural and reversal learning [25,28]. Individuals with ASD have decreased IGF1 in serum and CSF [29-31]. Together these studies provide insight into early IGF1 insufficiency from placenta as a possible mechanism of NDD-risk from preterm birth and adversity involving placental insufficiency. No published work has evaluated this mechanism of placental IGF1 insufficiency for neurodevelopmental risk [9,13].

In this study we aimed to determine whether placental *Igf1* insufficiency can drive translationally relevant outcomes, emphasizing the importance of the striatum. Placental *Igf1* insufficiency was induced using placental-targeted CRISPR manipulation [34] Significant changes in striatal development and forebrain white matter (WM) cellular composition relevant to NDD-risk were identified in males and females. These findings are critical as they identify possible therapeutic targets for impacts of perinatal neurodevelopmental risks including prematurity and placental insufficiency.

## Materials and Methods

### Animal Husbandry and Care

CD-1 mice purchased from Charles River or bred in house were used for CRISPR manipulation experiments. Tpbpαr/Adaf-AdaP-cre mice [35] provided by Dr. Yang Xia’s laboratory at Texas A&M Health Science Center were bred to create the placenta-cre line with Jax *Igf1* flox mice [36]. Sires were heterozygous Cre positive, while dams were Cre negative. Mouse handling and husbandry were performed as approved by the University of Iowa IACUC (protocol 3121284). Mice were housed in a 12-hour light cycle with food and water ad libitum.

### Placental-Targeted CRISPR Manipulation

Placental-targeted CRISPR manipulation was performed as previously described [34]. IGF-I CRISPR/Cas9 KO Plasmid (m) and Control CRISPR/Cas9 plasmids (Santa Cruz Biotechnology) were diluted to 0.073 µg/µl. Litters collected at embryonic timepoints had individual placentas injected with either IGF1 knockout or control plasmid, but litters used for postnatal assessments were treated with only one type of plasmid as previously described [13]. Placentas from experimental litters not directly manipulated are referred to as untreated [34].

### Sex Genotyping

Sex was determined for embryonic samples via PCR assessing *Jarid* or *Rmb31* genotype. Tpbpαr/Adaf-AdaP-cre/*Igf1* flox (i.e. placenta-cre) animals were genotyped using *Cre* and *Igf1* flox PCR primers (**Supplemental Table 1**).

### Quantitative Polymerase Chain Reaction

Placentas from CRISPR manipulated litters were collected as previously described [13]. Placentas from placenta-cre litters were cut in half laterally. One half had the decidua removed, the other retained the decidua to parse expression changes due to maternal influence. RNA was isolated using the Trizol method and concentrations were normalized prior to cDNA synthesis. PowerSYBR Master Mix was used for qPCR on the ViiA 7 Real-Time PCR System (Thermo Scientific). Gene expression was normalized to the housekeeping gene *18s*. See supplemental table 1 for primer sequences. The ddct method was used to generate normalized fold change from same-sex controls.

### ELISA

E14 placentas and embryo bodies (trunk, head removed) were digested for BCA and ELISA as previously described [13,34]. The BCA Protein Assay Kit (Thermo Fisher Scientific, 23227) and Mouse/Rat IGF-I/IGF-1 Quantikine ELISA Kit (R&D MG100) were performed per manufacturer instructions. Samples were run in duplicate. SkanIt Software (Thermo Scientific) was used to analyze results after absorbance values were measured from a plate reader.

### Embryonic Brain, Adult Brain, and Placental Histology

Embryonic day (E)14 and E18 heads were fixed, cryoprotected, and serially sectioned. Adult brains were collected approximately 2 weeks (12-14 postnatal weeks) after behavior completion via transcardial perfusion with saline followed by 4% PFA, cryoprotected, and serially sectioned. Staining was performed as previously described for NeuN, Olig2, and GFAP in adult brains [13]. Brain sections were mounted with Vectashield with DAPI (Vector Labs). Sections were visualized on an Axio Imager M.2 microscope (Zeiss) and StereoInvestigator (Microbrightfield) was used to quantify region volumes and unbiased stereology of cell populations.

Paraffin-embedded 5µm placental sections were H&E stained, visualized, and the area of placenta subregions were measured. E14 placenta sections were stained for GFP to demonstrate spatial incorporation of Igf1-KO CRISPR. Autofluorescence is displayed (**Supplemental Fig. 2C,D**) to distinguish placental subregions as previously reported [34]. Images were taken on an upright compound fluorescence microscope (Olympus). More details are available in **Supplemental Methods**.

### Forebrain RNA Sequencing

E18 forebrains were microdissected, stored in RNAlater, and processed for RNA using the RNAeasy Mini Kit (Qiagen). RNA concentrations were quantified using Qubit and quality was evaluated by Bioanalyzer. Only high-quality RNA (RIN >8) was used. RNA samples were sequenced on Illumina NovaSeq (Novogene). Illumina adapters were removed using TrimGalore (v 0.6.6), aligned to the mm10 reference transcriptome, then quantified using Salmon (v 1.10.1). Mapping efficiency was sufficient (**Supplemental Table 2**). Using R Studio, differentially expressed genes (DEGs) were identified using DESeq2 (v 1.46.0) [37]. All results were generated separately by sex and with litter as covariate. Using the svaseq function from the sva package (v 3.54.0) one surrogate variable was used for males and females to control for unwanted variation [38]. Significance was defined by a false discovery rate (FDR) of <0.05. DEGs were compared to the SFARI database list of autism-relevant genes from the 7/8/25 release list. Gene set enrichment analyses (GSEA) were performed for the DEGs identified from DESeq2 via the GSEA program (v4.4.0) to identify different enriched biological processes between treatment groups. Significance was defined as FDR<0.05 and trend as FDR<0.1.

### Maternal Behavior

Dams with manipulated litters were monitored as previously reported [13,39]. On postnatal day (P) 0, P3, P8, and P14 dams were recorded starting in the light cycle for 24, 3, 2, and 1 hours, respectively. Dams were analyzed in four-minute increments and scored by a blinded experimenter to assess maternal time on the nest.

### Behavioral Testing

Juvenile (P26-28) or early adult (P56+) mice were behaviorally tested as previously described [13]. All testing was done after habituation in home cages for a minimum of 30 minutes with task order as follows, least to most stressful.

### Open Field Testing (OFT)

Mice were individually placed inside a plexiglass OFT apparatus, recorded for 30 minutes, then analyzed using ANYmaze (Stoelting) to assess distance and center time in the first five minutes and throughout.

### Rotarod

Rotarod testing was performed for two consecutive days, 5 trials each day as previously described [13]. Weighted learning coefficients were then generated as the learning coefficient divided by the average of the first 2 trials. More details are available in **Supplemental Methods**.

### Y-Maze

Mice were placed in the same starting position in a Y-shaped apparatus (Stoelting), and arm entries were recorded for 5 minutes. Spontaneous alternation was calculated as proportion of alternating entry triplets to total arm entry triplets.

### Elevated Plus Maze (EPM)

Mice were placed into the center of the apparatus (Stoelting) and recorded for 5 minutes for assessment of open arm time using ANYmaze (Stoelting).

### Water T Maze

Water T maze consists of two phases: habit and reversal. Both phases of this task were performed as previously described [13]. More details are available in **Supplemental Methods**.

### Stereotypy

Mice were video recorded for 20 minutes in a rectangular plexiglass chamber. The number of stereotyped behaviors were scored by a blinded observer in 1-minute increments every 5 minutes.

### NDD-Relevant Behavior Composite Score

An NDD-relevant behavior composite score to compare degree of difference in behavior versus same-sex controls was generated for each mouse as previously described [10,13]. Each behavioral result was z-scored within each sex and the absolute values of z-scores were averaged. Included results were: OFT total distance, OFT total center time, Y-maze score, EPM open arm time, water T maze habit errors, water T maze reversal errors, and total stereotyped behaviors.

### Statistical Analysis

All statistical analysis was performed separately by sex for offspring outcomes. The ROUT test was used to remove outliers. The linear mixed effects model (lmer) with litter as a random effect in R was used for embryonic and placental analyses [13]. Postnatal and maternal measures were analyzed (Graphpad Prism) using Welch’s t-test or Mann-Whitney test (if F variance score failed) [13]. Placenta-cre analyses were run with either Welch’s t test or two-way ANOVA with post-hoc comparisons that did not include litter as a covariate due to a lack of within-litter pairings.

## Results

### Successful generation of placental Igf1 insufficiency model

Multiple methods were considered to reduce placental *Igf1*. Two placental Cre transgenic lines were considered: Cyp19-cre and Tpbpαr/Adaf-AdaP-cre. Cyp19-cre is expressed within a subset of trophoblasts, while Tpbpαr/Adaf-AdaP-cre is expressed in most trophoblasts [35,40], due to this difference Tpbpαr/Adaf-AdaP-cre was bred with an *Igf1* flox line (**Supplemental Fig. 1A**) [36]. Placental *Igf1* expression was not reduced in total placenta; only after isolation of the fetal zones to eliminate the unaffected source from maternal decidua cells was a decrease detected (**Supplemental Fig. 1B-D**). Body and placental mass were reduced regardless of Cre, suggesting undesirable impacts of *Igf1* flox alone (**Supplemental Fig. 1E-H**). No change was detected in embryonic forebrain development (**Supplemental Fig. 1I-L**). This model was not pursued due to the multiple limitations.

Placental-targeted CRISPR manipulation was used to reduce *Igf1* expression on E12 to assess whether insufficiency would influence placental, embryonic, and postnatal neurobehavioral outcomes (**Fig. 1A**). The *Igf1* knockout (Igf1-KO) CRISPR plasmid (**Fig. 1B**) successfully reduced placental *Igf1* across both sexes (**Fig. 1C and Supplemental Fig. 2A**) followed by a detectable decrease in IGF1 protein in the body of E14 males (**Supplemental Fig. 2B**). CRISPR incorporation was detected across placental subregions at E14 with the strongest staining in the decidua and junctional zone (**Supplemental Fig. 2C-F**). As previously reported, female placental *Igf1* was lower at baseline than males, which may influence response to manipulation (**Supplemental Fig. 2G**) [13]. Due to these findings, the placental-targeted CRISPR manipulation was the more effective method to induce placental *Igf1* insufficiency.

**Figure 1.**
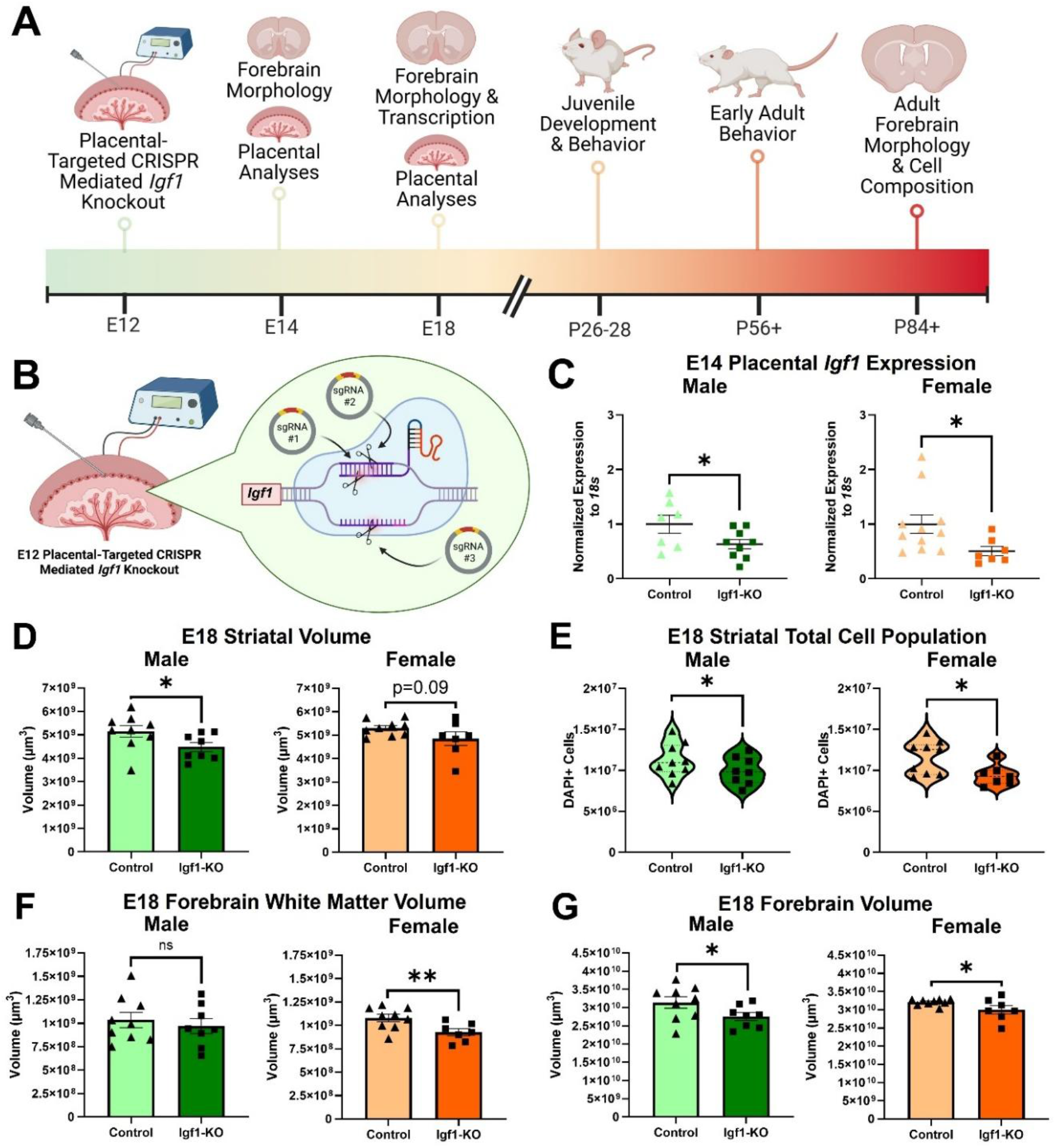
Successful generation of placental *Igf1* insufficiency model followed by impaired embryonic forebrain development. **(A)** Schematic of project timeline. **(B)** Schematic of E12 placental-targeted CRISPR mediated *Igf1* knockout. **(C)** Placental *Igf1* expression normalized to *18s* in E14 Igf1-KO males and females (n=7-11 per group). **(D)** E18 male and female striatal volume (n=7-9 per group). **(E)** E18 male and female striatal total cell (DAPI) population (n=7-9 per group). **(F)** E18 male and female forebrain white matter volume (n=7-9 per group). **(G)** E18 male and female forebrain volume (n=7-9 per group). All graphs show mean and SEM except for panel E which displays median and quartiles. ns=nonsignificant, trending p<0.1, and *p<0.05 by linear mixed effects model with litter as random effect.

There were few changes to placental structure and function from reduced placental *Igf1*, allowing this model to reproduce an insufficient supply of IGF1 to the fetus mimicking conditions like placental insufficiency. E14 Igf1-KO male placentas demonstrated minor compensations in IGF signaling expression that may promote IGF1 signaling, while females and E18 males did not (**Supplemental Fig. 2H,I**). Angiogenic factors were unchanged, IGF-dependent amino acid transporter expression was minimally altered, and placental subregions were unchanged (**Supplemental Fig. 2J-L**). Despite this, Igf1-KO females, but not males, had decreased placental and body mass at several timepoints, with body mass recovered by the juvenile timepoint (**Supplemental Fig. 3A-N**) supporting an insufficient supply of IGF1 for only prenatal growth.

### Placental Igf1 insufficiency impairs embryonic forebrain development

Forebrain morphology was assessed at E14 and E18 to evaluate impacts on early neurodevelopment. No changes in E14 ganglionic eminence and cortical volume or ganglionic eminence total cell population were identified, but an early decrease in forebrain volume was seen in females (**Supplemental Fig. 4A-E**). By E18, striatal volume and total cell population were reduced in both males and females (**Fig. 1D,E**). E18 females had decreased forebrain WM volume (**Fig. 1F**). There was decreased forebrain volume (**Fig. 1G**) and trend decreased cortical volume across sexes at E18 (**Supplemental Fig. 4F**), brain phenotypes relevant to preterm infant outcomes.

### Sex-specific ASD-relevant transcriptional changes in embryonic forebrains

E18 forebrain transcriptomic analysis identified changes relevant to NDDs, particularly ASD-risk (**Fig. 2A-C and Supplemental Fig. 5A,B**). The DEGs identified in Igf1-KO males and females compared to same-sex controls did not overlap, demonstrating clear sex-specific changes in E18 forebrain (**Supplemental Table 3 and 4**). Some key DEGs inform these different effects. In males, upregulation was found for *Cacna2d4*, a gene influencing neuromotor development, as the most significantly changed DEG [41]. In females, results showed the most significant DEG change was downregulation of *Huwe1*, a gene for a ubiquitin ligase that influences neuronal proliferation and differentiation [42].

**Figure 2.**
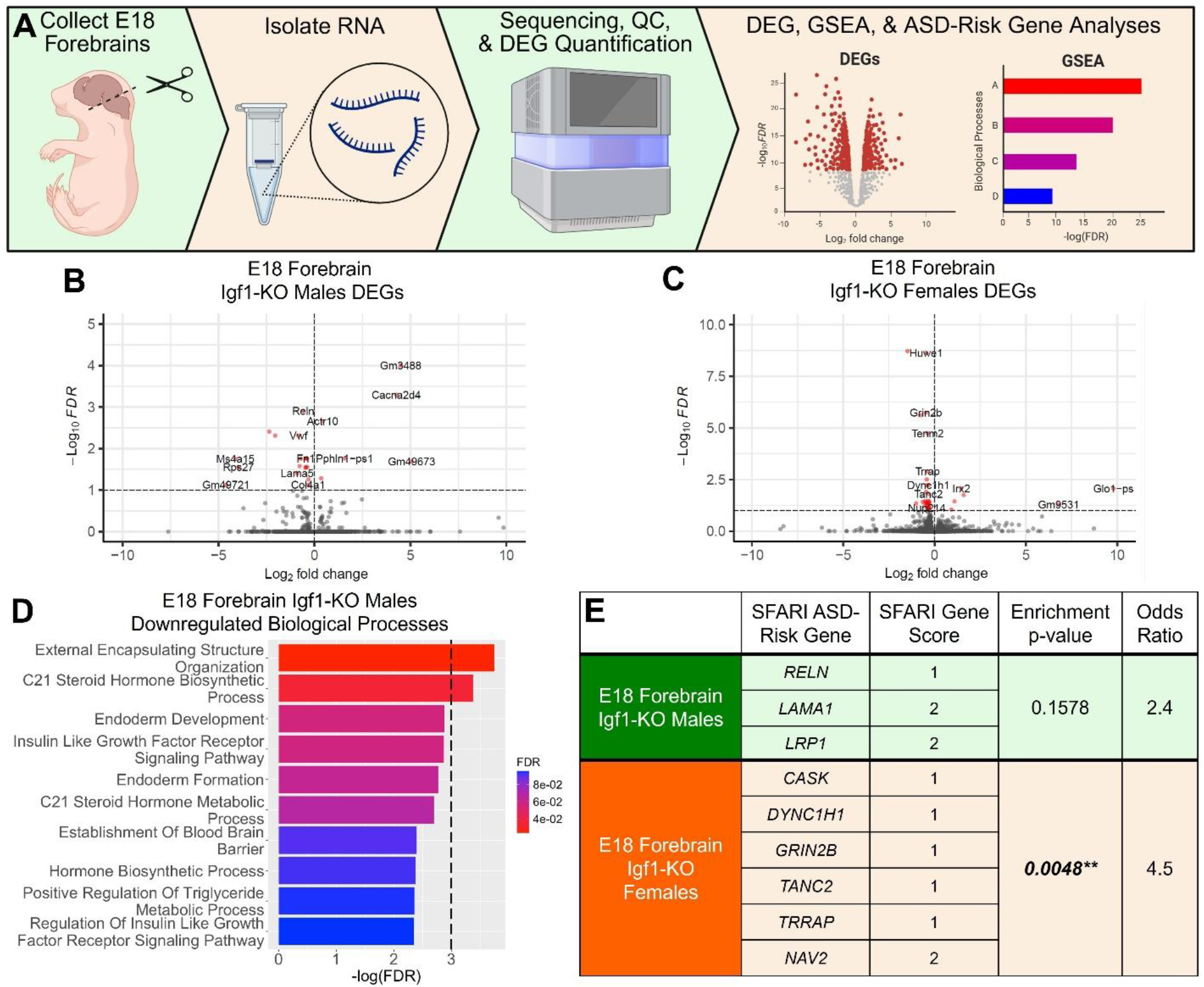
Sex-specific ASD-relevant transcriptional changes in embryonic forebrains. **(A)** Schematic of RNAsequencing workflow. Volcano plots showing DEGs from E18 Igf1-KO **(B)** male and **(C)** female forebrains compared to same-sex controls (n=6 per group). Scaling adjusted for readability, full volcano plots in supplement. **(D)** Key downregulated biological pathways in E18 Igf1-KO male forebrains identified using GSEA. Dashed line represents FDR = 0.05. Bars above this line (FDR<0.05) represent significant change while bar graphs below this line (FDR<0.1 but above 0.05) show trending change. **(E)** A table highlighting DEGs from E18 Igf1-KO male and female forebrains that are listed in the SFARI-ASD risk gene dataset (n=6 per group); significance defined as p<0.05.

GSEA analysis of biological processes did not identify changes in females. In contrast, Igf1-KO males had multiple downregulated relevant biological processes, including insulin-like growth factor receptor signaling (**Fig. 2D and Supplemental Table 5**). Igf1-KO male forebrain had unexpected downregulation of hormone biosynthesis, specifically C21 steroid hormones. This includes testosterone, which may influence the trajectory of sexually dimorphic neurodevelopmental processes [43,44]. There was downregulation of external encapsulating structure organization (i.e., extracellular matrix) and endoderm development and formation, involving differential expression of multiple laminin genes (*Lama1, Lama5, and Lamc1*) and *Reln*. Not only do these genes play critical roles in cell biology of the developing brain, but they are ASD-risk genes [45]. DEGs were assessed for enrichment for SFARI ASD-risk genes. While ASD-risk genes were not enriched in male forebrain DEGs, three were established ASD-risk genes (**Fig. 2E**). DEGs identified in Igf1-KO female forebrains were enriched for ASD-risk genes (**Fig. 2E**). Most of the ASD-risk genes found in females are implicated in synapse processes [46-51]. Both male and female DEGs that overlapped with SFARI risk genes were in the top two gene score groups indicating a high confidence association with ASD-risk (**Fig. 2E**).

### NDD-relevant behavioral deficits in juvenile Igf1-KO mice

We next examined how placental *Igf1* insufficiency influenced neurobehavior persistently into postnatal development. First, maternal care across postnatal timepoints was unchanged by the placental manipulations (**Supplemental Fig. 6A-F**), making further outcomes more likely to be due to altered placental *Igf1* directly.

To understand impacts on juveniles P26 to P28, motor activity, exploration, and motor learning were assessed. The first five minutes of locomotor activity was increased in Igf1-KO females and trend increased in males (**Fig. 3A,B**). Igf1-KO males had increased center time within this same period (**Fig. 3C**). These findings demonstrate an acute behavioral response to novelty, potentially due to increased stress susceptibility [52,53]. This was supported by no distance or center time differences across all 30 min of open field testing (**Fig. 3D,E**). Rotarod testing revealed Igf1-KO males improved less with motor learning, while females showed no deficit (**Fig. 3G,H**).

**Figure 3.**
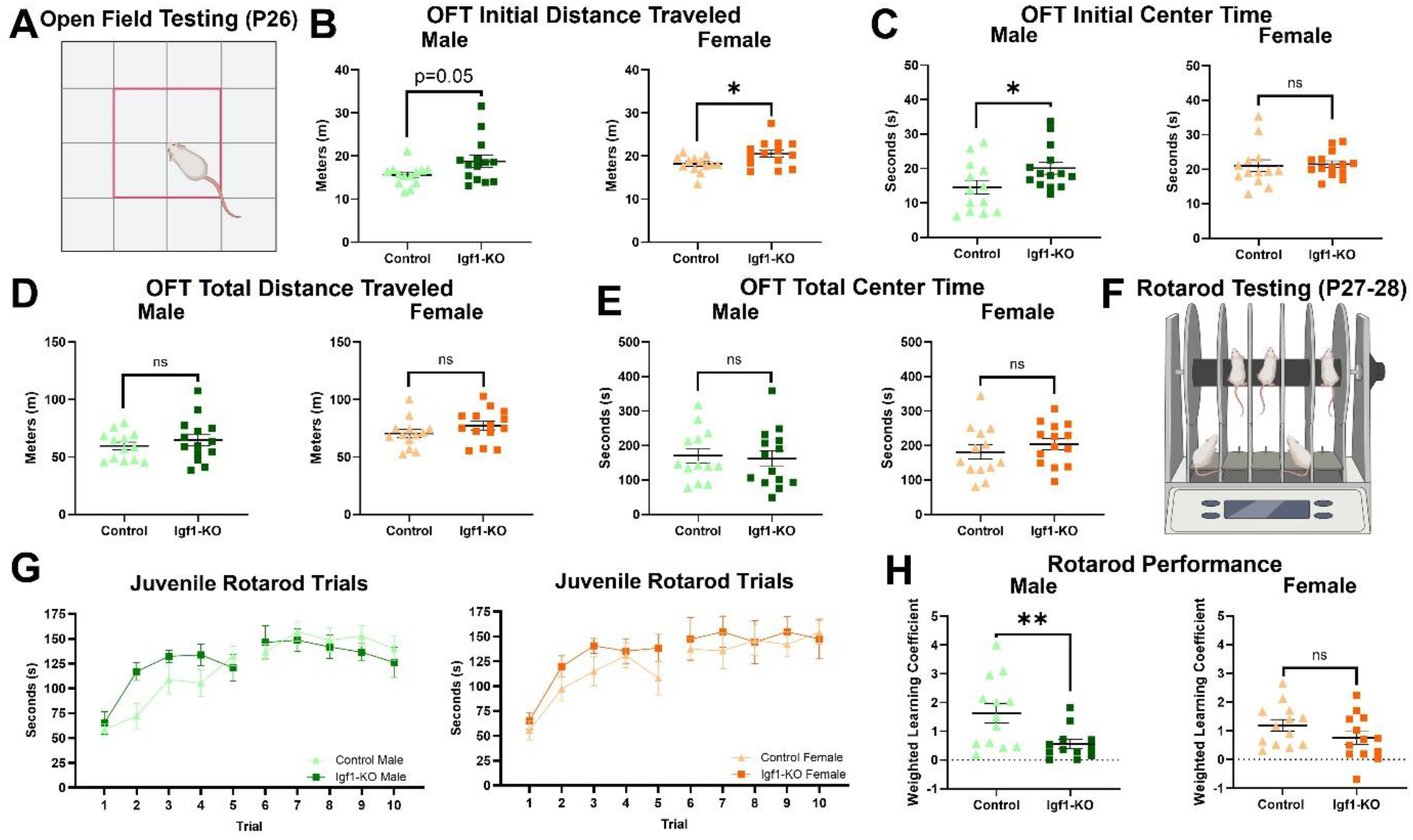
NDD-relevant behavioral deficits in juvenile Igf1-KO mice. **(A)** Representative graphic of open field testing (OFT). **(B)** Juvenile male and female distance traveled in the initial period (first 5 minutes) of OFT (n=13-14 per group). **(C)** Juvenile male and female center time in the initial period of OFT (n=13-14 per group). **(D)** Juvenile male and female distance traveled for the total time (30 minutes) of OFT (n=13-14 per group). **(E)** Juvenile male and female center time for the total time of OFT (n=13-14 per group). **(F)** Representative graphic of rotarod testing. **(G)** Juvenile male and female rotarod performance displayed as weighted learning coefficient (n=12-13 per group). **(H)** Graph of group means for each rotarod trial (n=12-13 per group). All graphs show mean and SEM. ns=nonsignificant, trending p<0.1, *p<0.05, and **p<0.01 by Welch’s t-test.

### Adult Igf1-KO males and females demonstrate opposite changes to NDD-relevant behavior

Behavior was assessed in early adulthood to determine lasting impacts relevant to striatal function and NDDs. Adult Igf1-KO showed more changes in striatal-dependent outcomes—habit/reversal learning and stereotyped behaviors—but less in domains such as working memory and anxiety-relevant behaviors (**Supplemental Fig. 7A-F**). While no differences in water T maze trials to habit learning criterion were identified in males or females across groups, Igf1-KO females had increased errors in habit learning and increased trials to reach reversal learning criterion (**Fig. 4A-C and Supplemental Fig. 7G**). Interestingly, Igf1-KO males performed better than controls on water T maze reversal learning (**Fig. 4C,D**). Stereotyped behavior analysis further identified opposite changes in Igf1-KO males and females, as males had increased stereotyped behaviors while females had a decrease (**Fig. 4E,F**). These differences suggest disrupted striatal development and/or function [25]. To compare behaviors across domains, NDD-relevant behavior composite scores were generated from adult behavioral tests as previously described [10,13] demonstrating distinct behavioral changes in Igf1-KO males and females (**Fig. 4G-J**). These results persist months after placental *Igf1* insufficiency, demonstrating a potential mechanism for NDD-relevant behavior after perinatal adversity.

**Figure 4.**
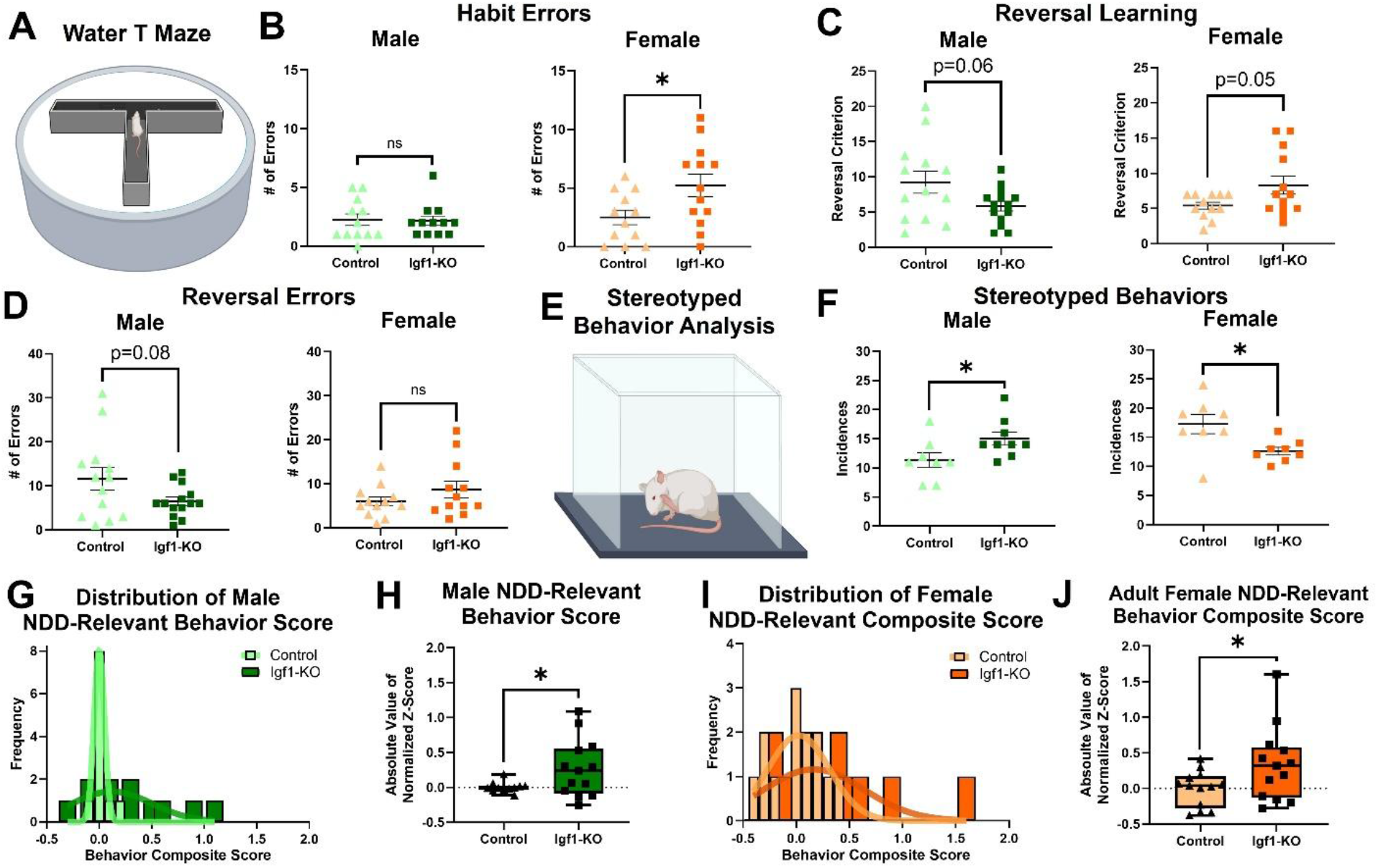
Adult Igf1-KO males and females demonstrate opposite changes to NDD-relevant behavior. **(A)** Representative graphic of the water T maze. **(B)** Number of errors made by adult male and females during water T maze habit training (n=12-13 per group). **(C)** Reversal learning in water T maze in adult male and females scored as reversal criterion (n=12-14 per group). **(D)** Number of errors made by adult male and females during water T maze reversal testing (n=12-14 per group). **(E)** Representative graphic of stereotyped behavior testing. **(F)** Incidences of adult male and female stereotyped behaviors (n=8-9 per group). **(G)** Distribution of adult male NDD-relevant behavior composite scores (n=11-13 per group). **(H)** Adult male NDD-relevant behavior composite score displayed as absolute value of normalized z-score (n=11-13 per group). **(I)** Distribution of adult female NDD-relevant behavior composite scores (n=13 per group). **(J)** Adult female NDD-relevant behavior composite score displayed as absolute value of normalized z-score (n=13 per group). All graphs show mean and SEM except panels H and J which show median and quartiles. ns=nonsignificant, trending p<0.1, and *p<0.05 by Welch’s t-test.

### Placental Igf1 insufficiency results in sex-specific persistent changes to the forebrain

The neurobiological underpinnings of behavioral changes were investigated by assessing adult forebrain. Similar to the placental *Igf1* overexpression study, striatal and cortical volumes that were affected embryonically no longer showed differences (**Fig. 5A-C** and **Supplemental Fig. 8A,B**) [13]. Despite this, Igf1-KO males had decreased striatal neurons, while females had a normalization (**Fig. 5D**). The sex-specific change to forebrain neuron population occurred in the context of sex differences in embryonic forebrain transcriptomic response to *Igf1* insufficiency which may represent the earliest stage in the trajectory of these effects. There was no difference in total striatal cell population in either sex (**Supplemental Fig. 8C**). A similar decrease of male prefrontal cortex neurons demonstrated that this effect affected multiple forebrain regions (**Supplemental Fig. 8D-F**). Due to embryonic changes in female forebrain WM volume, this structure was further investigated in the adult brain. Forebrain WM volume was undisrupted, (**Fig. 5E-H** and **Supplemental Fig. 8G**) but there was an increase in WM astrocytes in both males and females (**Fig. 5I-K**) while oligodendrocytes and total cell number was unchanged. This mirrors other models of preterm birth and placental insufficiency in which increased astrocytes is called reactive astrogliosis [33,54-57]. By contrast, no difference was found in striatal astrocytes (**Figure 5L**). These outcomes are highly relevant to NDD-risk after preterm birth and placental insufficiency [32,33].

**Figure 5.**
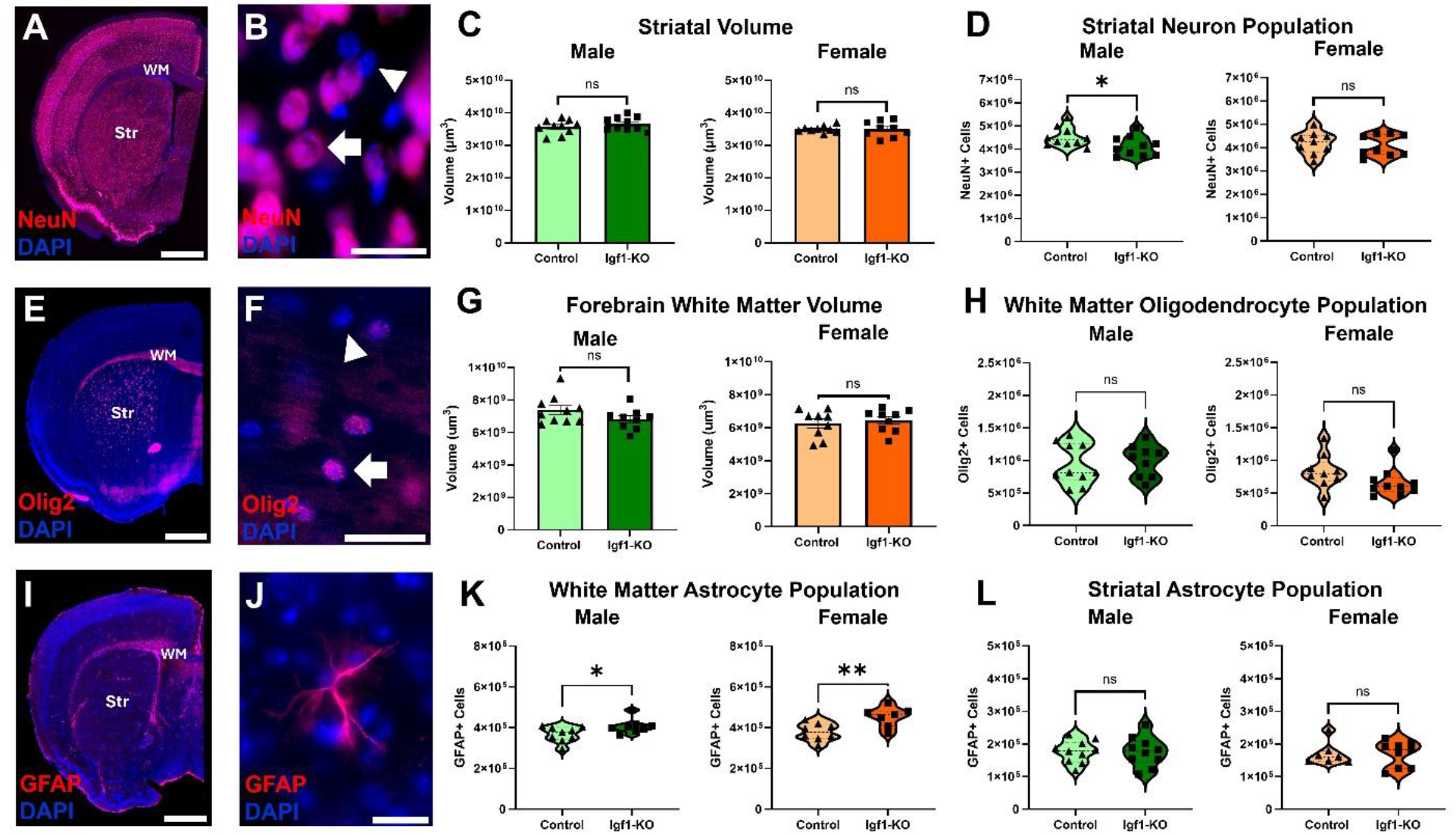
Placental *Igf1* insufficiency results in sex-specific persistent changes to the forebrain. **(A)** Representative image of a coronal hemisection of adult forebrain stained with DAPI (blue) and NeuN (red). Striatum (Str) and forebrain white matter (WM) labeled. **(B)** Representative image of DAPI (blue) and NeuN (red) staining in the adult striatum, arrow points to NeuN+ and DAPI+ cell and the arrowhead points to a DAPI+ cell. **(C)** Adult male and female striatal volume (n=9-10 per group). **(D)** Adult male and female striatal neuron (NeuN) population (n=9-10 per group). **(E)** Representative image of a coronal hemisection of adult forebrain stained with DAPI (blue) and Olig2 (red). Striatum (Str) and forebrain white matter (WM) labeled. **(F)** Representative image of DAPI (blue) and NeuN (red) staining in the adult forebrain, arrow points to Olig2+ and DAPI+ cell and the arrowhead points to a DAPI+ cell. **(G)** Adult male and female forebrain white matter volume (n=7-10 per group). **(H)** Adult male and female forebrain white matter oligodendrocyte (Olig2) population (n=9-10 per group). **(I)** Representative image of a coronal hemisection of adult forebrain stained with DAPI (blue) and GFAP (red). Striatum (Str) and forebrain white matter (WM) labeled. **(J)** Representative image of GFAP (red) staining in the adult forebrain. **(K)** Adult male and female forebrain white matter astrocyte (GFAP) population (n=7-9 per group). **(L)** Adult male and female striatal astrocyte (GFAP) population (n=7-10 per group). Scale bars display 1mm in panels A, E, and I, 25µm in panel B, and 20µm in panels F and J. Graphs in panels C and G show mean and SEM. Graphs in panels D, H, K and L show median and quartiles. ns=nonsignificant, *p<0.05, and **p <0.01 by Welch’s t-test.

## Discussion

This study identified that forebrain developmental and behavioral outcomes in mice relevant to NDDs can be driven by placental *Igf1* loss. Placental-targeted CRISPR manipulation allowed us to induce placental *Igf1*insufficiency that modeled hormone support reduction, in which production is diminished but not lost, similar to preterm birth, placental insufficiency, and other perinatal risks for NDDs [9,10,15-18,34]. Placental Igf1-KO mice had reduced embryonic forebrain growth, including specifically reduced striatal volume and cell density across sexes. However, sex-specific forebrain transcriptional changes suggested distinct developmental pathways in males and females alongside these overall structural effects. Following this, juvenile and adult behavior changes were distinct across sexes. The interaction between sex and early placental *Igf1* support of brain development was further emphasized by sex-specific changes in adult neuronal populations. Despite sex-specific effects, both adult males and females displayed increased forebrain WM astrocytes, similar to other perinatal adversity models which feature reactive astrogliosis [33,54-57]. Overall, placental *Igf1* insufficiency resulted in persistent effects on neurodevelopment and behavior that implicate this insufficiency as a possible mechanism of perinatal adversity contributing to NDD-risk.

Decreased embryonic forebrain and striatal volumes accompanied by decreased striatal total cell population in E18 Igf1-KO males and females showed a clear initial effect on forebrain development. This recapitulates human phenotypes that link low IGF1 levels with decreased brain volume after preterm birth [18]. Despite E18 striatal morphology not demonstrating sex-specificity, transcriptional changes in the E18 forebrain did not overlap in males and females. Females demonstrated enrichment of ASD-risk genes in embryonic forebrain DEGs which demonstrates one mechanism through which placental *Igf1* insufficiency may increase NDD-risk. Additionally, E18 Igf1-KO female forebrains demonstrated downregulation of *Huwe1*, a gene highly associated with intellectual disability and well-established as a major regulator of cell-cycle progression [42,58]. This downregulation emphasizes placental IGF1’s role in cell cycle regulation in neurodevelopment as similarly shown in a placental *Igf1* overexpression model [13]. Additionally, most of the ASD-risk genes identified in female forebrains are involved in synapse formation and function, key processes implicated in NDDs and ASD, highlighting another possible mechanism in that may increase NDD-risk [46-51]. While the striatum structurally normalized by adulthood, striatal-dependent behaviors were altered in females but in ways distinct from males, suggesting that the early distinct transcriptomics may have had lasting impacts. While both males and females had increased WM astrocytes, females had a more severe increase compared to their controls versus males. This follows our finding that embryonic forebrain WM volume was uniquely decreased in females.

In contrast, the influence of placental *Igf1* insufficiency on males resulted in early deficits in forebrain development which persisted. Early effects included a downregulation in the forebrain of IGF1R signaling, essential laminin genes, and hormone synthesis. While these results cannot be correlated with behavior, a decrease in perinatal hormone synthesis, particularly C21 steroids including testosterone and many others [59], may influence NDD-risk for males from this mechanism. Additionally, Igf1-KO males had reduced forebrain neurons which may contribute to downregulation of steroid hormone synthesis as neurons are primarily responsible for local hormone synthesis within the brain [59,60]. Interestingly, the behavioral changes displayed by males mostly resulted in behavioral phenotypes comparable to control females. This is somewhat in line with the general finding that female placentas express less *Igf1*, and this lower placenta *Igf1* may be one way that neurobehavioral outcomes are similarly shaped.

Both adult Igf1-KO males and females had a larger forebrain WM astrocyte population. This finding is significant as WM is one of the most common sites of brain injury after preterm birth [32,33]. Additionally, other models of perinatal adversity have similarly increased WM astrocytes; this suggests that the mechanism of placental *Igf1* insufficiency may influence such NDD-risk in these conditions [33,54-57,61-63]. Changes in WM astrocytes, including astrogliosis, are found in human preterm studies [33,64]. Increased GFAP and activated astrocytes in anterior cingulate cortex WM are found in children and adults with ASD [65-67]. Our model also has a persistent increase in WM astrocytes after placental *Igf1* insufficiency.

This study builds upon previous research demonstrating placental hormone insufficiency as critical mechanisms for NDD-risk. Similar to this study, placental allopregnanolone insufficiency in mice sex-specifically alters neurodevelopment and behavior relevant to NDDs [9,10,68]. Furthermore, a knockout of the placenta-specific *Igf2* transcript resulted in anxiety-like behavior and altered synaptic receptor expression in the hippocampus [69]. Lastly, our study builds upon a previous work that inhibited IGF1R activity in the early postnatal brain to model preterm birth in mice at a timepoint relevant to late *in utero* human brain development [70]. This study identified similar changes to neurodevelopment, including decreased prefrontal cortex neuron populations and increased stereotyped behaviors [70]. Our study expands upon this by isolating the role of placental *Igf1* on neurodevelopment.

While this study is highly informative, all brain regions and processes were not investigated, making it likely that other alterations occur that are relevant to placental *Igf1* insufficiency. Additionally, the timing of placental *Igf1* effects on neurodevelopment will invariably be different in mice than humans with perinatal adversity, as mouse brains are less developed than humans at birth. Despite these limitations, this study effectively demonstrates a clear role of placental *Igf1* in neurodevelopment that is informative for preterm birth and NDDs.

As preterm birth is associated with highly prevalent NDDs, it is crucial to understand how early loss of placental hormonal support impacts neurodevelopment. This study highlights a critical role of placental *Igf1* in neurodevelopment and its strong association with NDD-risk. The findings of this and similar studies emphasize the potential of early detection as well as intervention for placental hormone production anomalies in conditions such as preterm birth, intrauterine growth restriction, and placental insufficiency. This research will contribute to intervention development in infants at risk for NDDs and may aid in the creation of *in utero* treatments, as the placenta is an accessible target during prenatal development.

## Acknowledgements

The authors would like to acknowledge the following funding sources: National Institute of Mental Health grant R01MH122435 (HES), National Institute of General Medical Sciences grants T32GM008629 and T32GM145441 (AJC), National Center for Advancing Translation Science R25TR004393 (AG), Graduate College, University of Iowa (AJC), and Pappajohn Biomedical Institute, University of Iowa (FMF and BWQH). The authors would like to thank Dr. Yang Xia’s laboratory at Texas A&M Health Science Center for providing us with their Tpbpαr/Adaf-AdaP-cre mouse line. The authors would like to thank Dr. Val Sheffield and Dr. Calvin Carter’s labs at the University of Iowa for the use of their surgery room and equipment. We would also like to thank the Genomics Core at the University of Iowa. Figure 1A, figure 1B, figure 2A, figure 3A&F, figure 4A&E were created using BioRender.com Carver, A. (2026) https://BioRender.com/98ah0o5, https://BioRender.com/560kv8r, https://BioRender.com/3l1ahqd, https://BioRender.com/sgrxpq8, and https://BioRender.com/lie9v89.

## Author Contributions

Conceptualization, A.J.C. and H.E.S.; data curation, A.J.C and B.W.Q.H.; formal analysis, A.J.C., B.W.Q.H, and H.E.S.; funding acquisition A.J.C., F.M.F., B.W.Q.H, A.G., and H.E.S.; investigation A.J.C., F.M.F., R.J.T., B.W.Q.H., A.G., and R.T.F.; methodology, A.J.C., R.J.T., B.W.Q.H, and H.E.S.; project administration, A.J.C., R.J.T., and H.E.S.; resources, H.E.S.; supervision, A.J.C. and H.E.S.; validation, A.J.C. and R.J.T.; visualization, A.J.C., F.M.F., B.W.Q.H., A.G., and R.T.F.; writing—original draft, A.J.C., F.M.F., R.J.T., A.G., and R.T.F.; writing—review & editing, A.J.C., F.M.F., R.J.T., B.W.Q.H., and H.E.S.

## Data Availability

Upon publication, the transcriptomic data will be deposited in the Gene Expression Omnibus database. All other data will be available upon request.

