## Supplemental Material for "Placental Insulin-like Growth Factor 1 Insufficiency Drives Neurodevelopmental Disorder‑Relevant Behavioral Changes with Sex‑Specific Vulnerabilities"

### SUPPLEMENTAL INFORMATION

#### Supplemental Materials and Methods

Embryonic Brain, Adult Brain, and Placental Histology

Embryonic day (E)14 and E18 heads were fixed, cryoprotected, and serially sectioned at 25μm. Adult brains were collected approximately 2 weeks (12-14 postnatal weeks) after behavior completion via transcardial perfusion with saline followed by 4% PFA, cryoprotected, serially sectioned at 50μm, blocked with 10% horse serum, and immunostained in 10% horse serum/phosphate buffered saline with antibodies for NeuN (Cell Signaling, rabbit monoclonal, catalog #12943) (1:300), GFAP (Invitrogen, rabbit polyclonal, catalog #10019) (1:500), or Olig2 (Millipore Sigma, rabbit polyclonal, catalog #AB9610) (1:5000) followed by the appropriate secondary (Thermo Fischer, goat anti-rabbit, Alexa-Fluor 594, catalog #A-11012) (NeuN and GFAP 1:500. Olig2 1:1000) and mounted with Vectashield with DAPI (Vector Labs). Every $20^{th}$ embryonic and every $10^{th}$ adult section of each brain was visualized on an Axio Imager M.2 microscope (Zeiss) and StereoInvestigator (Microbrightfield) was used to quantify region volumes and unbiased stereology of cell populations using consistent counting frames and randomly placed grids in the “optical fractionator” workflow.

Paraffin-embedded 5μm placental sections were H&E stained, then visualized on the Axio Imager M.2 microscope (Zeiss). The area of placenta subregions was measured using StereoInvestigator (Microbrightfield) from two sections and averaged. E14 placenta sections were stained for GFP (enQuire BioReagents, chicken polyclonal, catalog #QAB10251) (1:500) followed by the appropriate secondary (Thermo Fischer, goat anti-chicken, Alexa-Fluor 594, catalog #A-11042) (1:500) to demonstrate spatial incorporation of Igf1-KO CRISPR. Autofluorescence is displayed in green (**Supplemental Fig. 2C,D**) to distinguish placental subregions as previously reported [34]. Images taken on an upright compound fluorescence microscope (Olympus).

Rotarod

Rotarod testing was performed for two consecutive days, 5 trials each day as previously described [13]. Mice were placed on the apparatus (Ugo Basile) and accelerated from 4 to 80 rpm over 240 seconds. Holding onto the rod for two full revolutions or falling concluded the trial; duration recorded. Learning coefficients were calculated as the difference of the average trial length of the last 2 trials (4 and 5 of day two) and average trial length of the first 2 trials (1 and 2 from day one). Weighted learning coefficients were then generated as the learning coefficient divided by the average of the first 2 trials.

Water T Maze

A T-shaped maze was placed in a tub filled with water mixed with nontoxic white paint to obfuscate a platform inside one arm beneath the water's surface. Water T maze consists of two phases: habit and reversal. For the habit phase, mice were trained to find the hidden platform for 10 trials per day until training criterion was met: 5 consecutive trials with no incorrect arm entry (i.e. error). Training continued for 2 more days after criterion. For the reversal phase, the platform was moved to the opposite arm and trials proceeded only until criterion was met. Errors were recorded until both criteria were met. Habit and reversal learning performance were assessed by number of errors and trials before criterion, correcting trial number for lead-in trials on each day.

**Supplemental Table 1: Primer/oligonucleotide sequences.** List of primer/oligonucleotide sequences used in this paper for genotyping or qPCR.

**Supplemental Table 2. RNAsequencing mapping rate.** Table with mapping rate (listed as percent) to the mm10 genome for each E18 forebrain sample used for RNAsequencing.

**Supplemental Table 3. Full list of DEGs identified in E18 male Igf1-KO forebrain RNAsequencing.** Table with full list of DEGs identified in Igf1-KO males versus same-sex controls.

**Supplemental Table 4. Full list of DEGs identified in E18 female Igf1-KO forebrain RNAsequencing.** Table with full list of DEGs identified in Igf1-KO females versus same-sex controls.

**Supplemental Table 5. Extended list of downregulated biological processes identified in E18 male Igf1-KO forebrains.** All significantly and trending downregulated biological pathways in E18 Igf1-KO male forebrains identified using GSEA. Significant change was defined as FDR<0.05 and a trending change was FDR<0.1. Igf1-KO females had no significant or trending pathways, so data is not shown. Sample size was 6 per group.

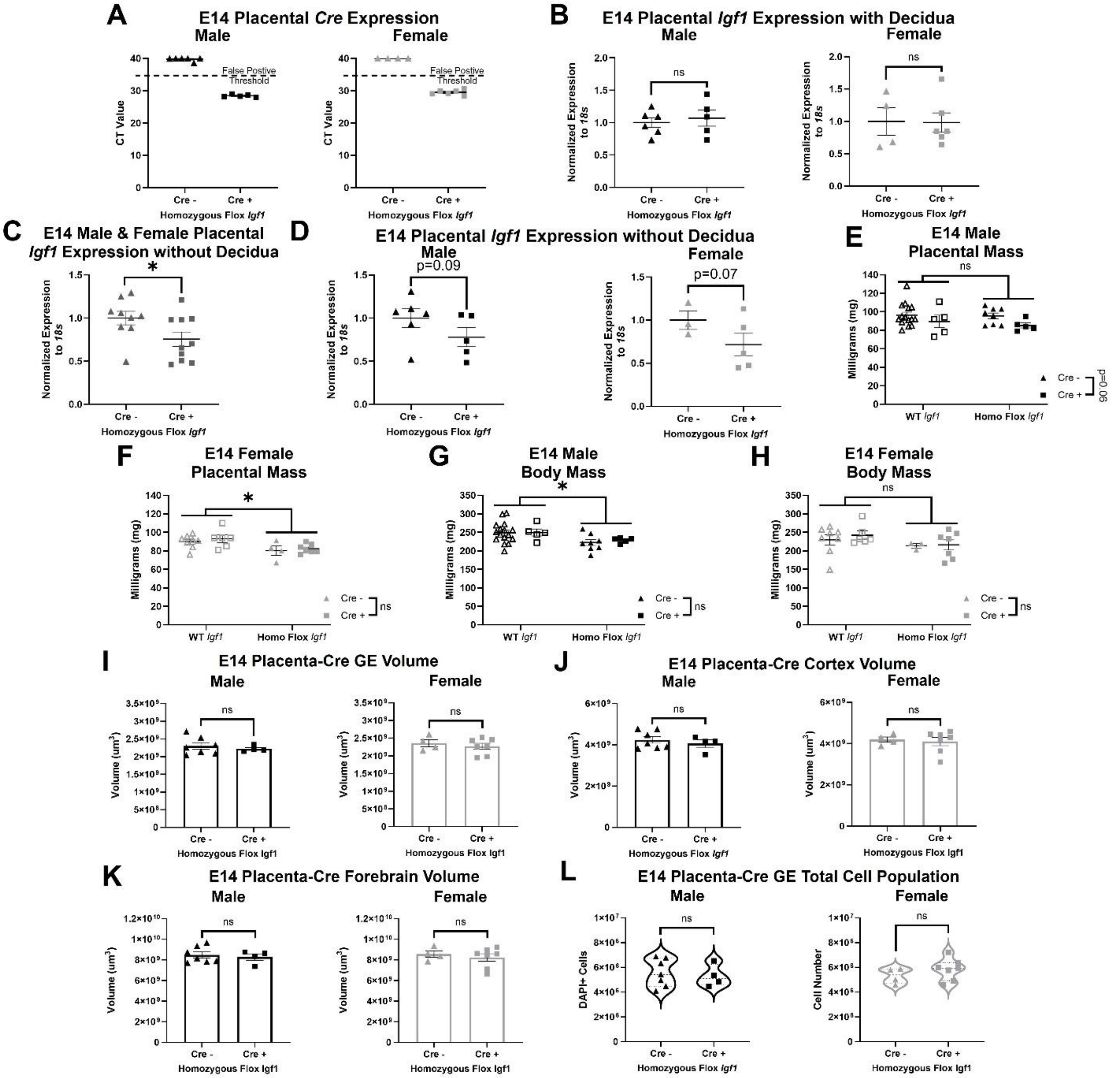


**Supplemental Figure 1. Findings from the placenta-cre line. (A)** E14 placenta-cre male and female cre expression shown as CT value in homozygous flox animals (n=4-6 per group). **(B)** Expression of placental *Igf1* normalized to *18s* from placentas with decidua attached in placenta-cre males and females (n=4-6 per group). Expression of placental *Igf1* normalized to *18s* from placenta-cre placentas with the decidua removed in **(C)** males and females combined and **(D)** males and females separately in homozygous flox animals (n=3-10 per group). E14 placental mass in placenta-cre animals *Igf1* wildtype (WT) and homozygous flox (Homo Flox *Igf1*) **(E)** males and **(F)** females (n=4-15 per group). E14 body mass in placenta-cre animals *Igf1* wildtype (WT) and homozygous flox (Homo Flox *Igf1*) **(G)** males and **(H)** females (n=3-16 per group). **(I)** E14 placenta-cre male and female ganglionic eminence (GE) volume (n=4-7 per group). **(J)** E14 placenta-cre male and female cortex volume (n=4-7 per group). **(K)** E14 placenta-cre male and female forebrain volume (n=4-7 per group). **(L)** E14 placenta-cre male and female GE total cell (DAPI) population (n=4-7 per group). All graphs show mean and SEM except for panel L which displays median and quartiles. ns=nonsignificant, trending $p<0.1$, and *$p < 0.05$ by Welch's t-test for panels A-D and I-L, and by two-way ANOVA with multiple comparisons for panels E-H.

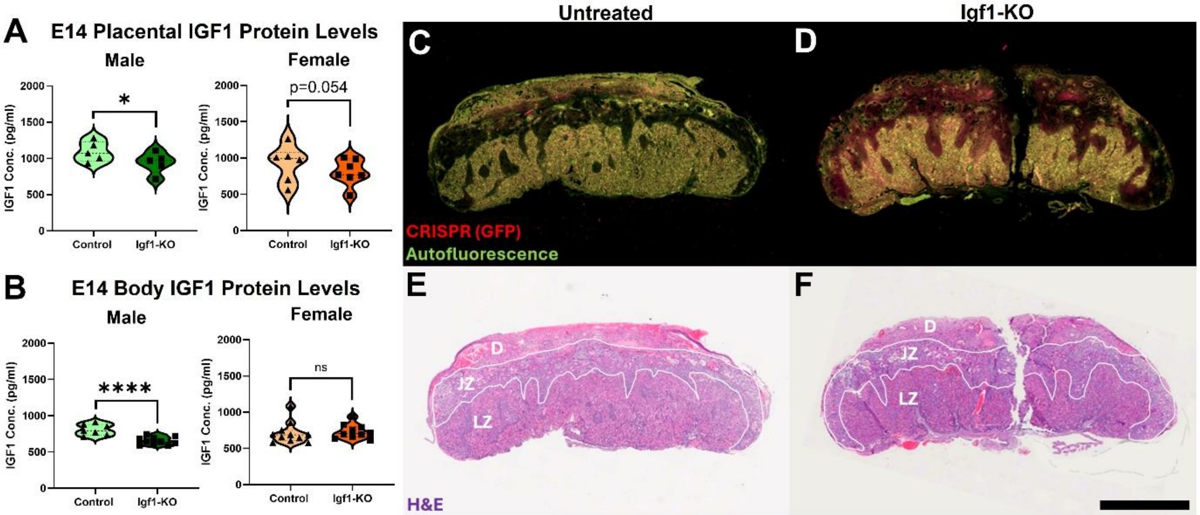


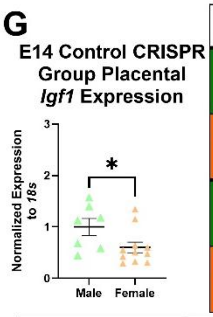


| H | *Igf2* | *Igf1R* | *Igf2R* | *Igfbp3* | *Igfbp5* |
|---|---|---|---|---|---|
| E14 Igf1-KO Males | ↑p= 0.078# (n=7-10) | p=0.798 (n=6-10) | ↓p=0.06# (n=5-8) | ↑p=0.035* (n=7-10) | p=0.221 (n=5-10) |
| E14 Igf1-KO Females | p=0.498 (n=9-11) | p=0.259 (n=7-9) | p=0.7287 (n=9-10) | p=0.999 (n=9-11) | p=0.713 (n=9-11) |
| E18 Igf1-KO Males | p=0.282 (n=9) | p=0.644 (n=9) | p=0.395 (n=9) | p=0.9 (n=9) | p=0.563 (n=7-9) |
| E18 Igf1-KO Females | ↓p=0.005** (n=9) | p=0.75 (n=9) | p=0.395 (n=9) | p=0.533 (n=9) | p=0.773 (n=8) |

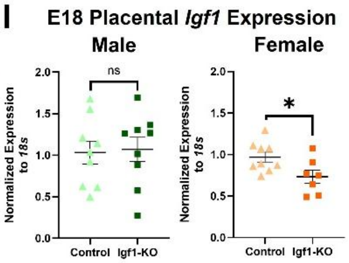


| J | *Plgf* | *Vegf* | *Flt1* | *Snat1* | *Snat2* | *Snat4* |
|---|---|---|---|---|---|---|
| E14 Igf1-KO Males | p=0.135 (n=7-10) | p=0.213 (n=7-10) | p=0.551 (n=7-10) | p=0.458 (n=6-10) | ↓p=0.081# (n=6-10) | p=0.213 (n=7-10) |
| E14 Igf1-KO Females | p=0.646 (n=5-8) | p=0.3869 (n=9-11) | p=0.568 (n=7-8) | p=0.306 (n=9-11) | p=0.242 (n=9-11) | p=0.893 (n=9-11) |
| E18 Igf1-KO Males | p=0.476 (n=9) | p=0.381 (n=9) | p=0.5 (n=9) | p=0.593 (n=9) | p=0.225 (n=9) | p=0.976 (n=9) |
| E18 Igf1-KO Females | p=0.103 (n=9) | p=0.954 (n=8-9) | p=0.715 (n=9) | p=0.366 (n=9) | ↓p=0.099# (n=9) | ↑p=0.032* (n=8-9) |

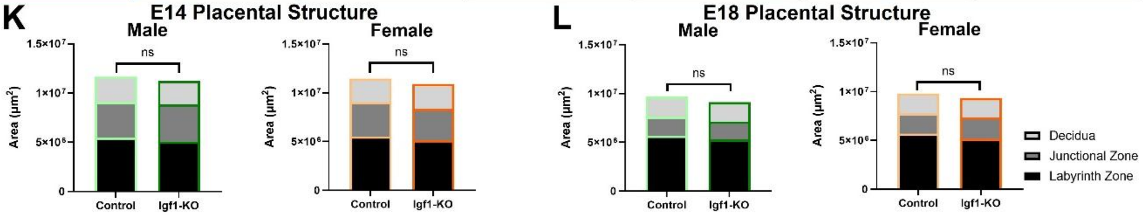


**Supplemental Figure 2. Placental assessments. (A)** E14 placental IGF1 protein levels in males and females (n=5-7 per group). **(B)** E14 body IGF1 protein levels in males and females (n=7-11 per group). E14 placenta **(C)** untreated and **(D)** Igf1-KO sections stained for CRISPR incorporation (GFP stained in red) and autofluorescence (green). H&E sections of E14 **(E)** untreated and **(F)** Igf1-KO placentas to highlight placental subregions. Placental subregions are labeled as decidua (D), junctional zone (JZ), and labyrinth zone (LZ). Panels C and D are sister sections of the same placenta from panels E and F, respectively. **(G)** Placental *Igf1* expression normalized to *18s* in CRISPR control males and females (n=7-11 per group). **(H)** Table displaying placental expression of IGF signaling factors normalized to *18s* compared to age and sex-specific controls. **(I)** E18 placental *Igf1* expression normalized to *18s* in males and females versus same-sex controls (n=7-9 per group). **(J)** Table displaying placental expression of angiogenic and growth promoting factors normalized to *18s* compared to age and sex-specific controls. **(K)** E14 placental subregion area in males and females versus same-sex controls (n=5-9 per group). **(E)** E18 placental subregion area in males and females versus same-sex controls (n=7-9 per group). Panels A and C show mean and SEM. p-value and sample size listed within panels B, D, and G. Direction of change from control group labeled as arrows in panels B, D, and G. ns=nonsignificant, trending $p<0.1$, *$p < 0.05$, and **$p< 0.01$ by linear mixed effects model with litter as random effect.

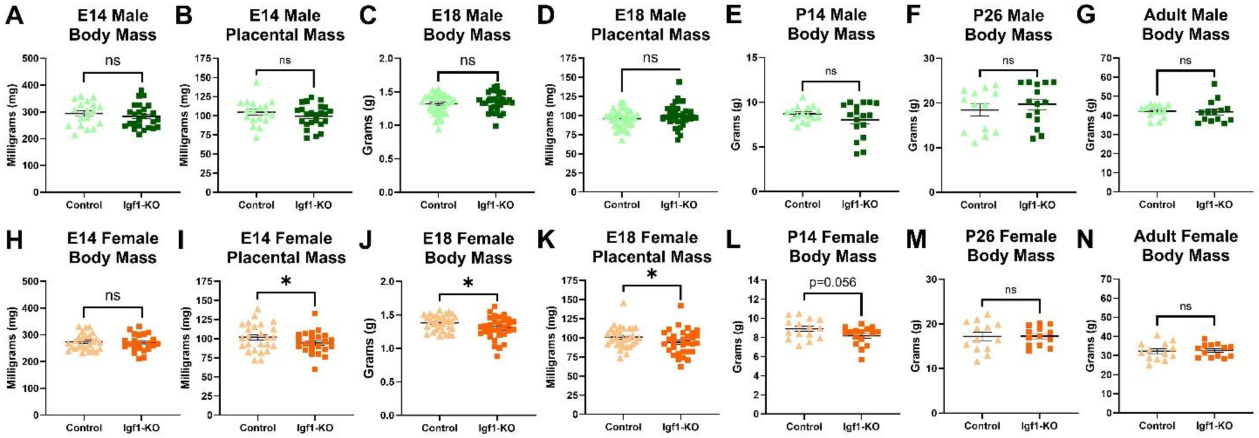


**Supplemental Figure 3. Placental and body masses. (A)** E14 body mass in males (n=18-25 per group). **(B)** E14 placental mass in males (n=18-24 per group). **(C)** E18 body mass in males (n=35-36 per group). **(D)** E18 placental mass in males (n=35 per group). **(E)** P14 body mass in males (n=16-17 per group). **(F)** P26 body mass in males (n=13-15 per group). **(G)** Adult body mass in males (n=13-14 per group). **(H)** E14 body mass in females (n=22-26 per group). **(I)** E14 placental mass in females (n=23-25 per group). **(J)** E18 body mass in females (n=31 per group). **(K)** E18 placental mass in females (n=31 per group). **(L)** P14 body mass in females (n=15-16 per group). **(M)** P26 body mass in females (n=13 per group). **(N)** Adult body mass in females (n=13 per group). All graphs show mean and SEM. ns=nonsignificant, trending $p<0.1$, *$p<0.05$ by linear mixed effects model with litter as random effect for embryonic comparisons and Welch's t-test for postnatal comparisons.

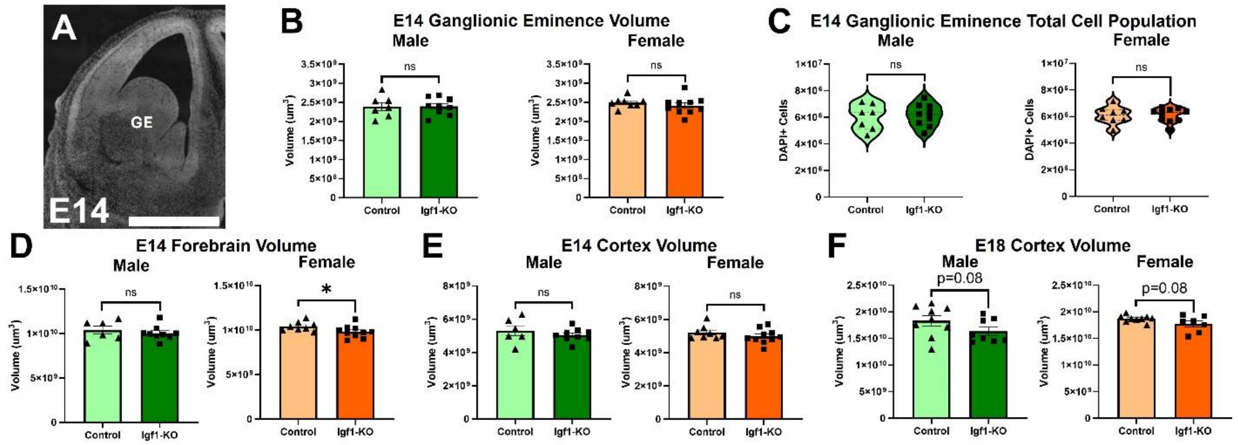


**Supplemental Figure 4. Further embryonic brain assessments after Igf1-KO. (A)** Representative image of a coronal hemisection of E14 forebrain stained with DAPI, ganglionic eminence (GE) labeled. **(B)** E14 male and female ganglionic eminence volume (n=7-10 per group). **(C)** E14 male and female GE total cell (DAPI) population (n=7-10 per group). **(D)** E14 male and female forebrain volume (n=6-10 per group). **(E)** E14 control and Igf1-KO male and female cortex volume (n=6-10 per group). **(F)** E18 control and Igf1-KO male and female cortex volume (n= per group). Scale bar displays 1mm in panels A. All graphs show mean and SEM except for panel C which displays median and quartiles. ns=nonsignificant, trending $p<0.1$, *$p<0.05$ by linear mixed effects model with litter as random effect.

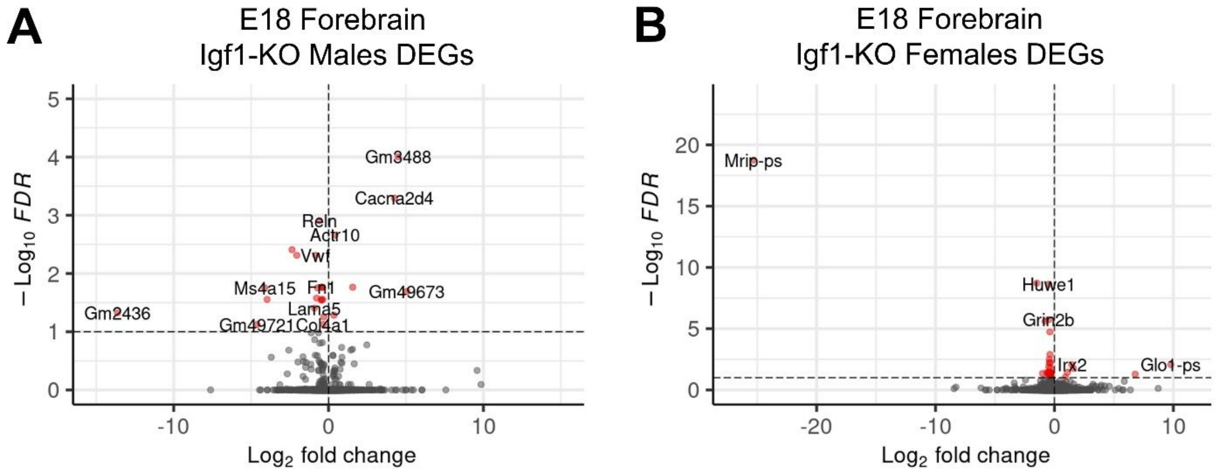


**Supplemental Figure 5. RNAsequencing volcano plots.** Volcano plots showing DEGs from E18 Igf1-KO **(A)** male and **(B)** female forebrains compared to same-sex controls (n=6 per group). Significance was defined as FDR<0.05.

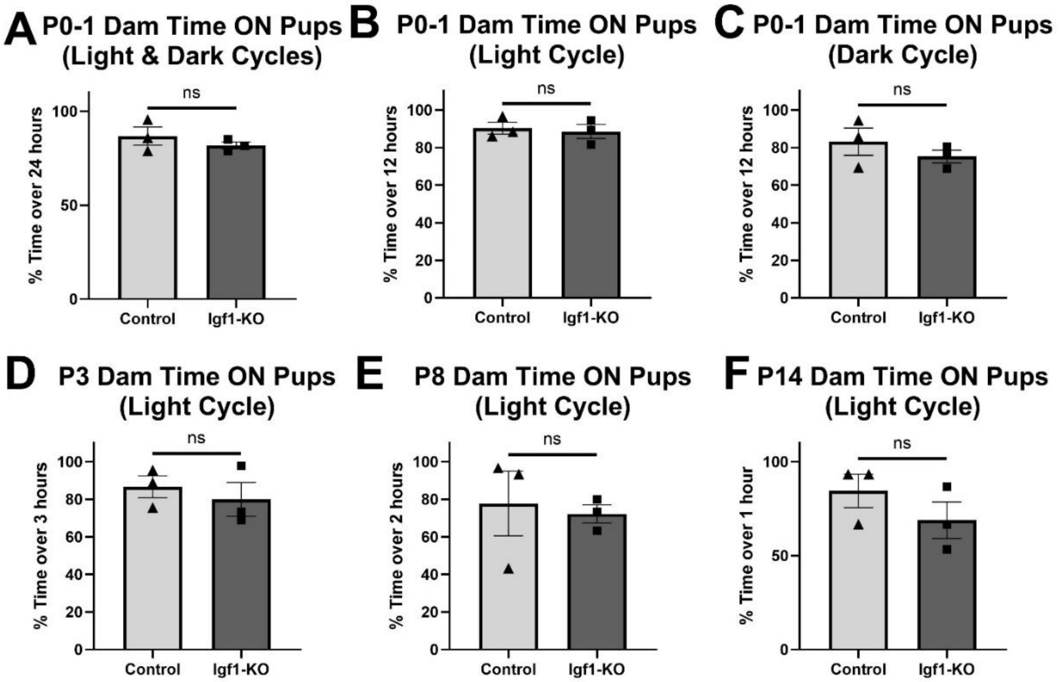


**Supplemental Figure 6. Maternal care after placental-targeted CRISPR manipulation. (A)** Dam time on pups during light and dark cycle for 24 hours from P0 to P1 (n=3 per group). **(B)** Dam time on pups during light cycle for 12 hours from P0 to P1 (n=3 per group). **(C)** Dam time on pups during dark cycle for 12 hours from P0 to P1 (n=3 per group). **(D)** Dam time on pups during light cycle for 3 hours on P3 (n=3 per group). **(E)** Dam time on pups during light cycle for 2 hours on P8 (n=3 per group). **(F)** Dam time on pups during light cycle for 1 hour on P14 (n=3 per group). All graphs show mean and SEM. ns=nonsignificant by Welch's t-test or Mann-Whitney test.

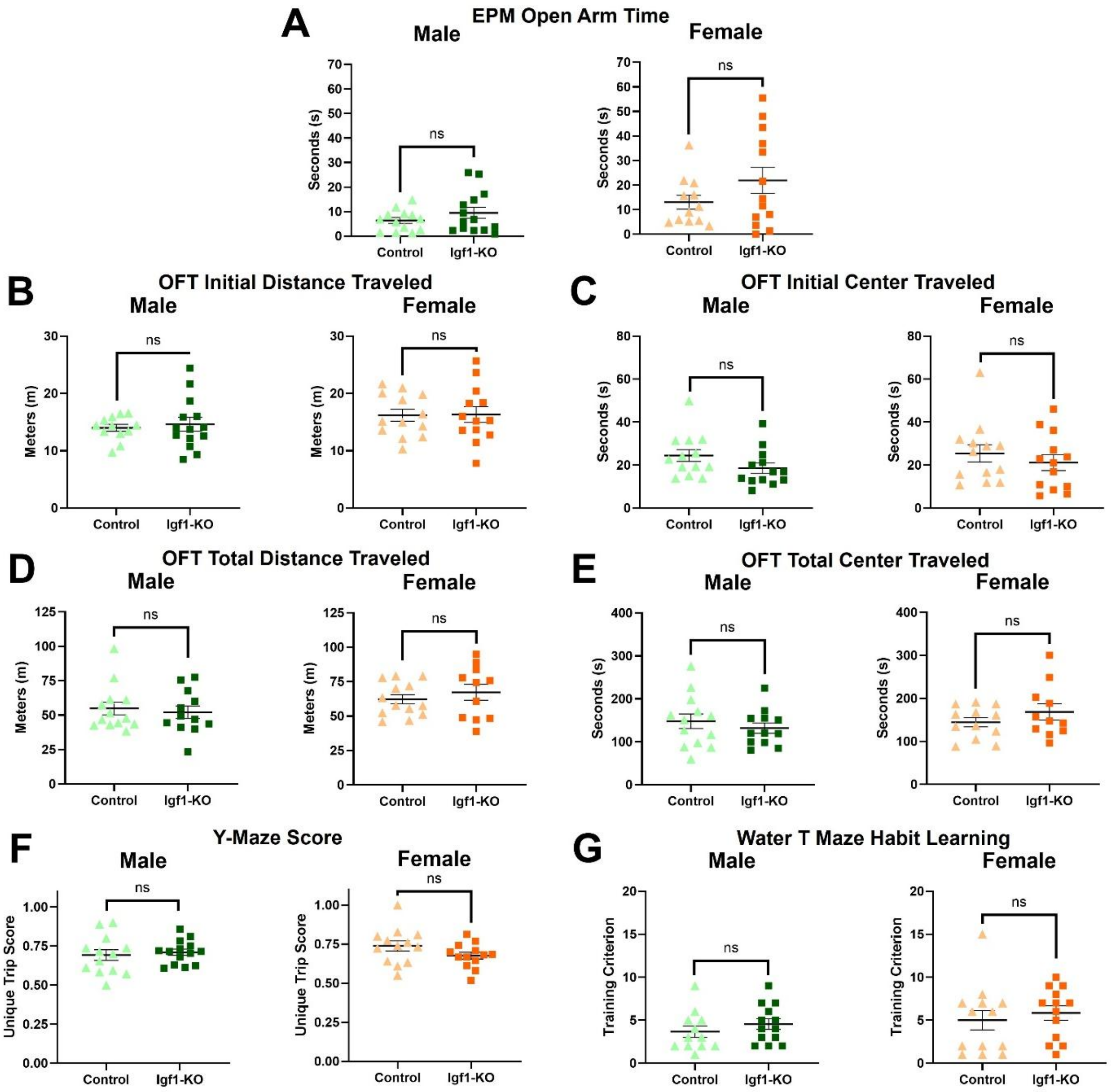


**Supplemental Figure 7. Adult elevated plus maze, open field testing, and Y-maze behavior findings. (A)** Adult male and female elevated plus maze (EPM) time spent in open arms (n=12-14 per group). **(B)** Adult male and female distance traveled in the initial period (first 5 minutes) of OFT (n=12-14 per group). **(C)** Adult male and female center time in the initial period of OFT (n=13 per group). **(D)** Adult male and female distance traveled for the total time (30 minutes) of OFT (n=11-13 per group). **(E)** Adult male and female center time for the total time of OFT (n=11-13 per group). **(F)** Adult male and female Y-maze unique trip score (n=12-14 per group). **(G)** Habit learning in water T maze in adult male and females scored as training criterion (n=12-13 per group). All graphs show mean and SEM. ns=nonsignificant by Welch's t-test.

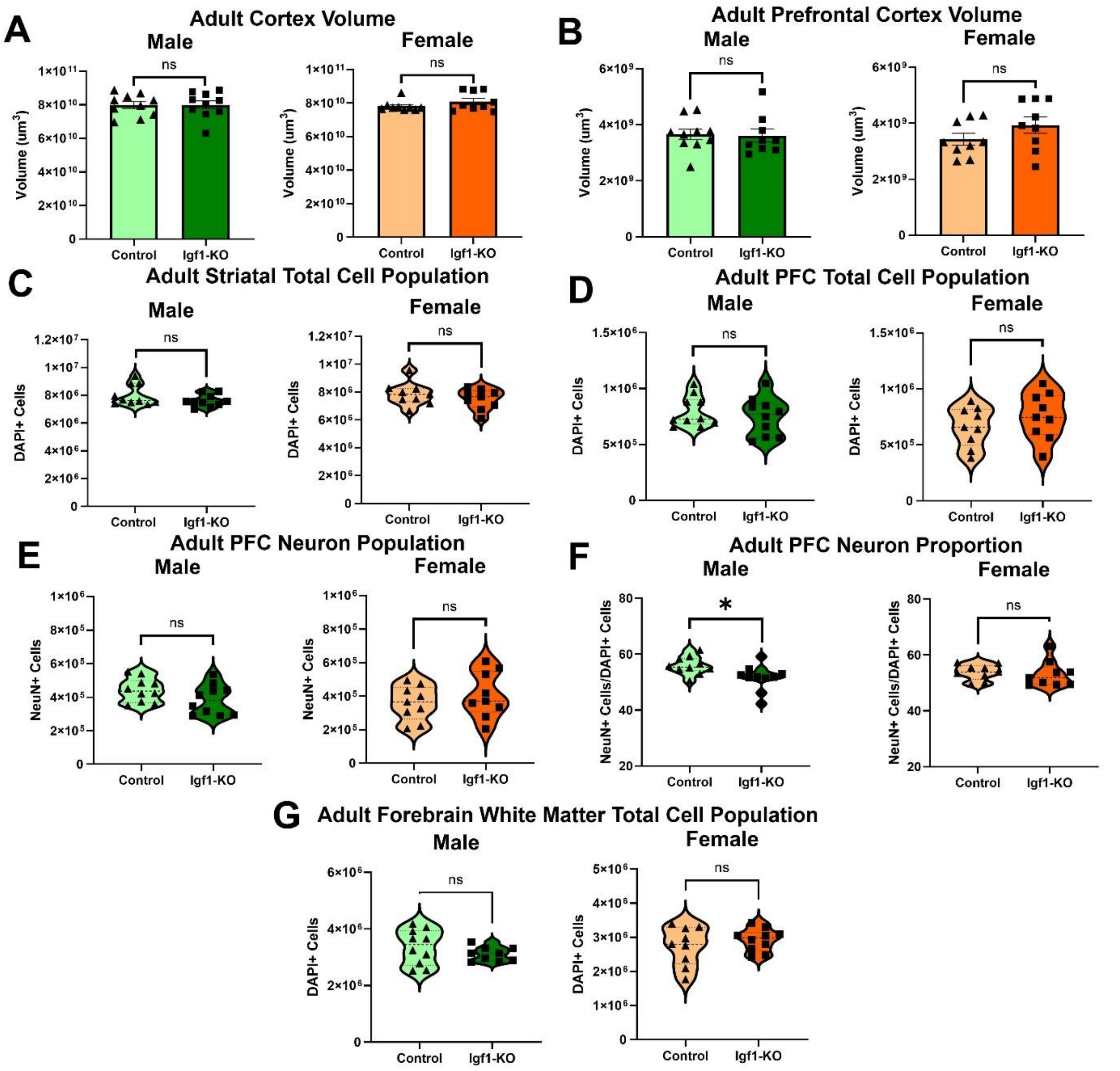


**Supplemental Figure 8. Further adult forebrain findings. (A)** Adult male and female cortex volume (n=9-10 per group). **(B)** Adult male and female prefrontal cortex (PFC) volume (n=9-10 per group). **(C)** Adult male and female striatal total cell (DAPI) population (n=9-10 per group). **(D)** Adult male and female PFC total cell (DAPI) population (n=9-10 per group). **(E)** Adult male and female PFC neuron (NeuN) population (n=9-10 per group). **(F)** Adult male and female PFC neuron proportion (NeuN/DAPI cells) (n=9-10 per group). **(G)** Adult male and female forebrain white matter total cell (DAPI) population (n=9-10 per group). Graphs in panels C and D show mean and SEM. Graphs in panels A, B, E, F and G show median and quartiles. ns=nonsignificant and *p<0.05 by Welch's t-test.
